# Lactose-based surfactants complexable with oligonucleic acids as gene delivery agents

**DOI:** 10.1101/2022.06.17.496588

**Authors:** Michalina Wilkowska, Monika Makrocka-Rydzyk, Katarzyna Michocka, Ryszard Zieliński, Krzysztof Sobczak, Maciej Kozak

## Abstract

Advances in molecular medicine related to the treatment of genetic disorders and cancer require finding new pathways for gene transfer. Nonviral delivery methods rely on the application of polymers, liposomes and cationic lipid systems used as vehicle. Among these systems, there is increasing interest in surfactants, which, due to their ability to complex with nucleic acids can deliver into cell DNA or RNA molecules of almost any size, which is unattainable with viral gene delivery systems.The main aim of this study was to determine the effect of the concentration of lactose-based surfactants (zwitterionic derivatives of sulfobetaine with carbohydrate moieties) on the structure of DNA/RNA as well as to explore their abilities of nucleic acid complexation. Structural studies of DNA or RNA in complexes with surfactants of two types at various concentrations were conducted using circular dichroism (CD) spectroscopy, gel electrophoresis (GE) and synchrotron radiation small angle X-ray scattering (SR-SAXS). Our studies showed that the examined surfactants have excellent properties of forming complexes with DNA and RNA. Additionally, to determine the cytotoxicity and transfection abilities of the studied lipoplexes, preliminary tests were performed in HeLa and fibroblast cells. The obtained results suggest that these systems have relatively low toxicity; however, further research is needed in this area.

## Introduction

Over 60 years have passed since the discovery of DNA structure; however, interest in nucleic acids has not diminished. Although built of relatively simple monomers, they take the form of many diverse complex spatial structures and have many functions, starting from coding of genetic information to enzymatic roles (ribozymes). Recently, considerable attention has been given to the possibilities of using nucleic acids as therapeutic substances, their delivery to cells and their practical medical applications [1]. A particularly interesting area of study was the potential use of all types of molecules selectively interacting with fragments of DNA within chromosomes or shorter sections of other nucleic acids [2,3]. Studies of the abovementioned issues have indicated the importance of knowing the conditions in which certain structures and spatial organizations of DNA or RNA are preserved, both in vitro and in vivo, which may be useful for the development of new solutions in anticancer or gene therapies. Knowledge of physical factors and chemical compounds determining the stability of proper structures of nucleic acids is expected to permit the development of new diagnostic methods at the molecular level and improve gene therapy methods. Interesting agents promoting the formation of condensed forms of nucleic acids are zwitterionic surfactants (having an additional saccharide group) capable of forming complexes with nucleic acids. Due to their specific spatial structure, these complexes of surfactants with nucleic acids (so-called lipoplexes) can act as carriers that are able to deliver DNA or RNA molecules of practically any size to the cells, and importantly, they are safe for living organisms [4,5]. It should be emphasized that intensive research work is currently focused on the antisense therapeutics. Antisense reagents are defined as short single-stranded oligonucleotides of ribonucleic acid (ssRNA) or deoxyribonucleic acid (ssDNA) or mixture of both that can bind to specific region within different classes of RNA, mostly mRNA, or genomic DNA. Depends on the structure and chemical modifications of bases or phosphodiester bonds of antisense reagents they can control expression of targeted genes/RNA on different levels of their metabolism, e.g. can induce degradation of mutant mRNA via different mechanisms. These molecules can be used in the therapies of certain viral diseases, cancers, autoimmunological diseases, cardiovascular problems or neurological diseases [2]. An important issue is development of safe system for delivery of antisense therapeutics to the cells of living organisms [1,6,7].

This study provides an extension of our previous preliminary investigations on complexation of one lactose-based surfactant with siRNA [8]. The main aim of this study was to determine the effect of increasing concentration of lactose-based zwitterionic surfactant in aqueous solution on the conformation of short single- and double-stranded nucleic acids and to determine the structure of these lipoplexes. The research was performed for two types of lactose-based zwitterionic surfactants differing in the length of alkyl chain, and among the nucleic acids under study were small interfering RNA duplexes (siRNA), single-stranded DNA (ssDNA) and double-stranded DNA (dsDNA).

The structural studies were performed using circular dichroism (CD), electrophoretic mobility shift assay (EMSA) in agarose gel and synchrotron radiation small angle X-ray diffraction method (SR-SAXS). Our objectives included also determination of cytotoxicity of studied surfactants and transfection properties of solution of these surfactants to verify the possibility of their use as a carrier of nucleic acids in molecular biology or gene therapy. The subject of our interest was the effect of complexation of surfactants with antisense ssDNA builded of 23 nucleotides sequence complementary to CUG repeats in RNA. Antisense oligonucleotides with this sequence were previously shown as gene therapy tools with high therapeutic potential in cellular and animal models of myotonic dystrophy (DM) [9,10,11]. As already mentioned, the systems of a surfactant and nucleic acid can be used for introduction of certain nucleic acids (DNA or RNA) to the cells. This process may lead to inhibition of expression of certain genes (gene silencing) usually with the use of siRNA [12]. Hence, we also investigated the effect of different conditions of lactose surfactants complexation with a short siRNA consisting of sequence of 21 base pairs. The sequence studied is responsible for silencing of *CELF1* gene, which expression is affected in myotonic dystrophy type 1 (DM1) [13–15]. Preliminary transfection tests using a selected fibroblast line were performed for a fluorescently labelled single DNA strand (ssDNA). Moreover, the toxicity of the lipoplexes and complex of lipolex with ssDNA and their transfection abilities towards two types of cells, fibroblasts and epithelial cells, with mutant form of genes related to the DM were characterized.

## Materials and methods

Two surfactants showing unique properties were chosen for the study: N-(3-propanesulfonate)-N-dodecyl-lactoside ammonium hydrochloride (LA12) and N-(3-propanesulfonate)-N-tetradecyl-lactoside ammonium hydrochloride (LA14). The LA12 surfactant has been previously examined by us but only in combination with siRNA [8], while this study is devoted to its application in the complexation of dsDNA and ssDNA. The surfactants were synthesized on the basis of the sugar group lactose – the hydrophilic part of the surfactant, linked through a nitrogen atom to a 1,3-propanesulfonate group and an alkyl chain (dodecyl or tetradecyl). The chemical structures of the surfactants studied are shown in Fig 1.

**Fig 1.**
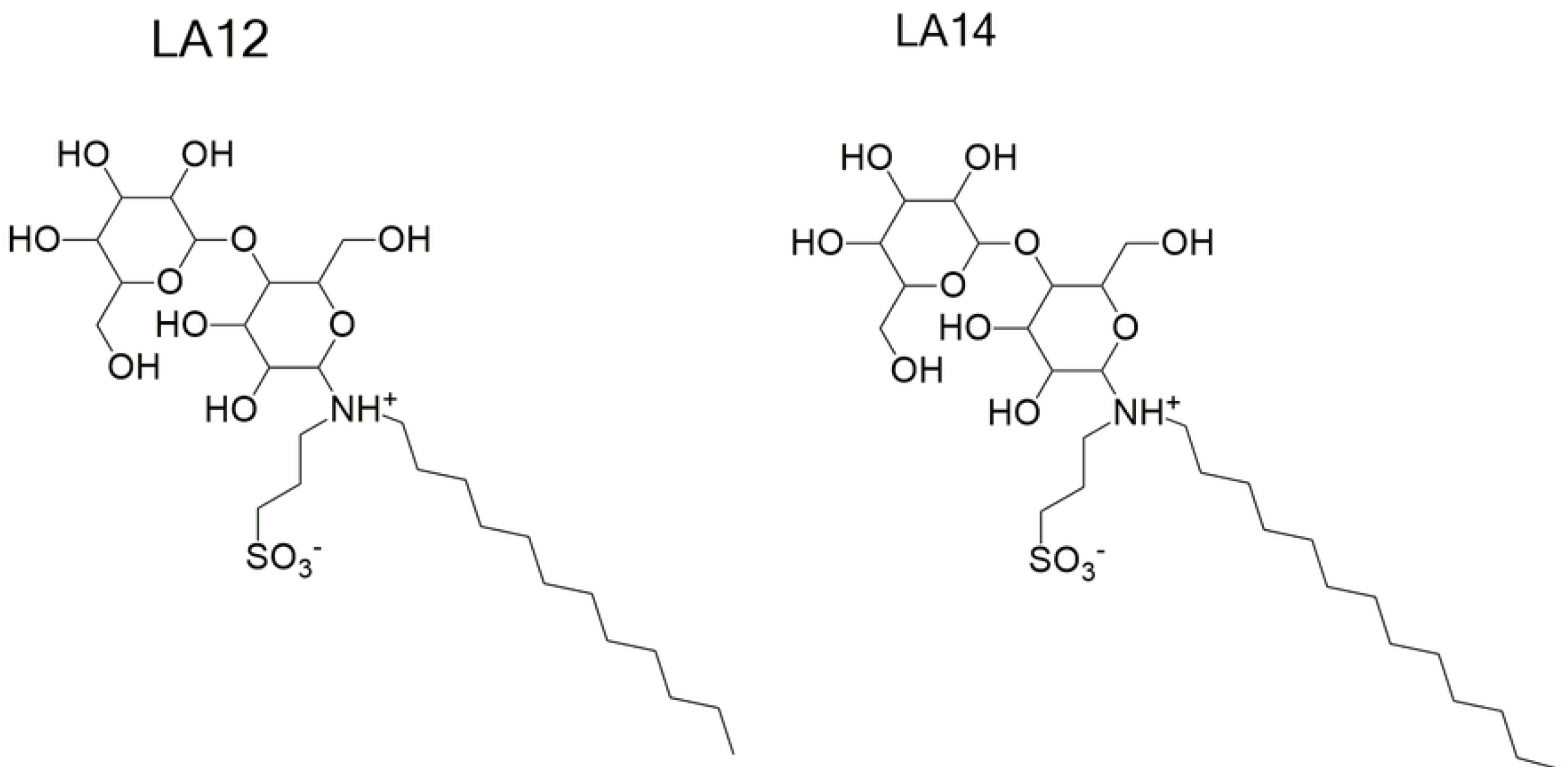
Chemical structures of N-(3-propanesulfonate)-N-dodecyl-lactoside ammonium hydrochloride (LA12) (left) and N-(3-propanesulfonate)-N-tetradecyl-lactoside ammonium hydrochloride (LA14) (right).

The three types of nucleic acids used in the study (i.e., siRNA, dsDNA and ssDNA) were synthesized by Future Synthesis (Poznań, Poland) and used without additional purification. The oligonucleotide sequences of the nucleic acids mentioned are given in Table 1.

**Table 1.**
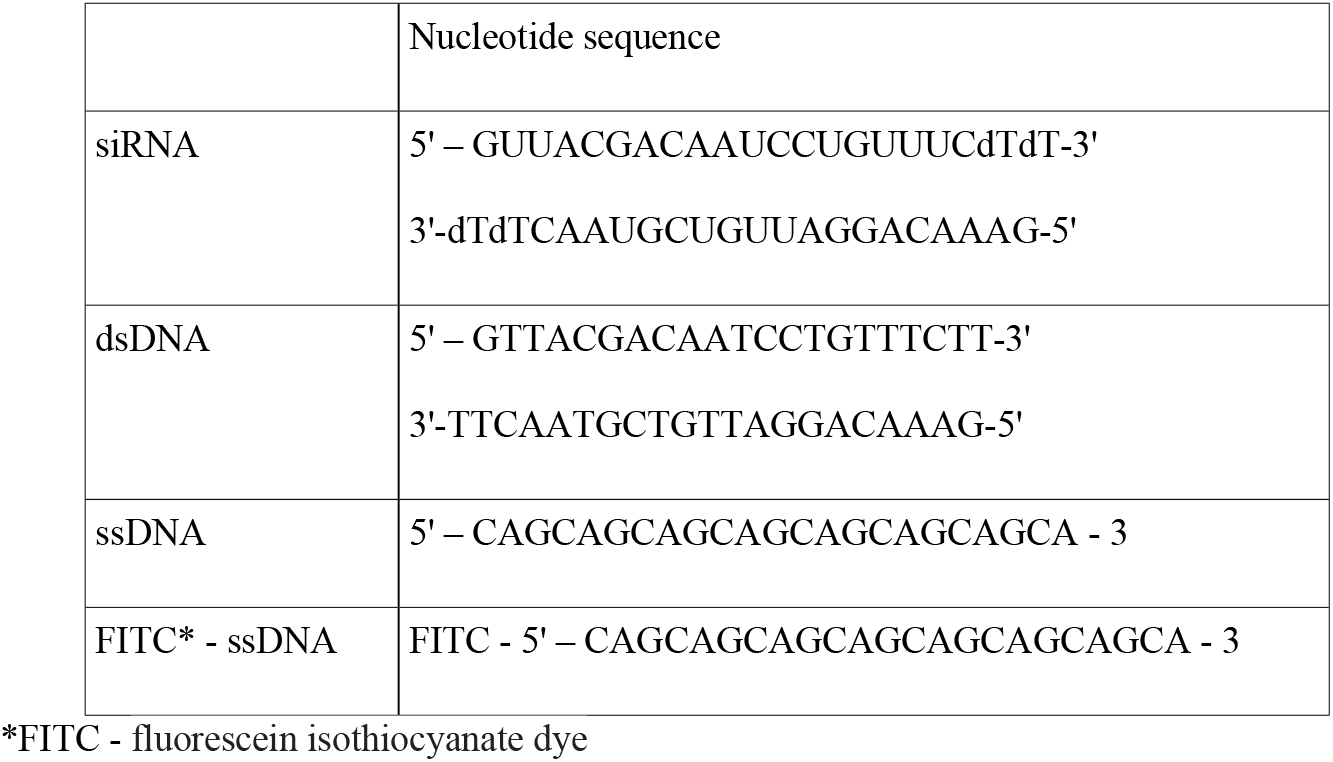
Oligonucleotide sequences of the nucleic acids used in the study.

The sequence of short double-stranded siRNA oligomer (21 base pairs) chosen for the study is responsible for silencing the genes targeted in the therapy of myotonic dystrophy of type 1 [13–15]. It has characteristic overhangs built of deoxythymidines to improve loading efficiency of this RNA interference to RISC complex and to increase its stability. The dsDNA sequence was chosen to be identical to the sequence of siRNA. We had already successfully used these two sequences for testing the carriers of nucleic acids based on gemini surfactants [16–20]. The short sequence of single-stranded DNA (23 nucleotides) was designed as a reference for potential studies on transfection efficiency [21,22]. Preliminary transfection studies concerning FITC-ssDNA were performed in this work.

### Sample preparation

All samples to be studied were prepared in 10 mM phosphate buffer (Na_2_HPO_4_NaH_2_PO_4_) of pH 7.03, using ddH_2_O, in nuclease-free medium. Lyophilized samples of the nucleic acids studied were dissolved directly in a water solution of the buffer and left to stand for at least 4 h prior to measurements at 4 °C. The surfactants dissolved in the buffer solution were incubated for 30 min at a temperature not exceeding 30 °C with mild sonication and then incubated for 15 min at room temperature. The procedure was repeated 5 times to achieve complete dissolution and homogenization of the solution. The ready-to-use surfactant solutions were kept at 4 °C. Prior to measurements, they were subjected to an additional sonication for 15 min at a temperature not exceeding 30 °C to break the aggregations that may be formed due to self-assembly processes. In solutions of surfactant and nucleic acid, the varied parameter was the surfactant concentration, while that of nucleic acid remained unchanged. To facilitate comparison of results, the fraction of surfactants relative to nucleic acid was expressed as the ratio of positive to negative charges (p/n). The values of p/n were calculated from the formula:

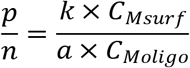

where *C_M surf_* and *C_M oligo_* denote the molar concentrations of the surfactant and nucleic acid, respectively. Symbol *k* represents the amount of the positive charge of the head of the surfactant molecule, while symbol *a* stands for the amount of the negative charge in the molecule of nucleic acid. Each lipoplex solution was prepared by mixing surfactant solution and appropriate nucleic acid solution at a ratio of 1:1 for 15 min at ambient temperature. Measurements were made for surfactant concentrations ranging from 1 mM to 160 mM, while the final concentration of siRNA and dsDNA during complexation was 47 μM, and for ssDNA 100 μM.

### Small angle X-ray scattering (SAXS)

SAXS studies of the studied lipoplex solutions were performed using the BioSAXS beamline P12, operated by EMBL, located at the Petra-III storage ring (DESY – Hamburg, Germany) [23]. All SAXS measurements were made with synchrotron radiation using an online sample-robot BioSAXS [24] with a thermostated capillary measuring chamber and a Pilatus 2M detector. The range of SAXS data collected corresponds to the scattering vector s range from 0.22 to 4.8 nm^-1^ (s=4πsin(θ)/λ, θ – scattering angle, λ – synchrotron radiation wavelength) [25]. All SAXS data were processed and analysed by the PRIMUS program [26].

### Circular dichroism (CD)

Conformations of the studied nucleic acid and lipoplex solutions were determined by circular dichroism using a Jasco J-815 spectropolarimeter (Jasco Inc., Japan). The spectral range of CD measurements was 200-320 nm. The measurements were made at room temperature with the samples placed in a quartz cuvette (0.5 mm beam path) with the following parameters: standard sensitivity, continuous mode, data integration time of 1 s, data pitch of 0.5 nm, scanning rate of 50 nm/min, and accumulation number of 5.

### Agarose gel electrophoresis (AGE)

Complexation of selected surfactants with nucleic acids was performed with electrophoresis in a 2% agarose gel matrix. The tests were performed in the presence of 0.5 μg/ml ethidium bromide and 1x TBE buffer (89 mM Tris – 89 mM boric acid – 2 mM EDTA, pH 8.3). The reference samples were pure oligonucleotide and DirectLoad™ Wide Range DNA Marker (Sigma–Aldrich). The samples of 8 μl in volume, subjected to complexation for 15 min, were deposited onto the agarose gel. The electrophoretic separation was performed at 120 V for 45 min. After separation, the gels were analysed in a transilluminator operated at λ= 300 nm.

### Cell culture

The cytotoxicity of the surfactants was examined by qualitative and quantitative methods towards two types of adherent cells differing in the sensitivity to external agents: epithelial HeLa cells and primary fibroblasts. Two lines of fibroblasts were used, GM04033 and GM07492.The following cell lines were obtained from the NIGMS Human Genetic Cell Repository at the Coriell Institute for Medical Research. The former line has a *DMPK* gene with expansion of 1000 repetitions of the triplets (CTG)n/(CAG)n in the 3’UTR (untranslated region). This gene is responsible for myotonic dystrophy type 1 (DM1) [21]. The second fibroblast line, GM07492, is the control line with a small number of the triplet repetitions in the *DMPK* gene. The fibroblasts were grown in complete EMEM medium (Lonza, catalogue number BE12-611F), containing 10% FBS (fetal bovine serum), 1% of antibiotic (antimycotic agents) and 1% of a mixture of amino acids. For HeLa cells culture, a complete DMEM medium (Lonza, catalogue number BE12-604F), containing 10% FBS and 1% antibiotic/antimycotic was used. Cells were grown at 37°C in the presence of 5% CO_2_.

### Qualitative assessment of surfactant toxicity – microscopic observations

For determination of cytotoxicity, the cultivation medium was changed to incomplete medium, without FBS or antibiotics. The cytotoxicity tests were performed in 12-well plates in 1 ml of cultivation medium. To each well containing the medium with the cells, a surfactant solution was added, and the contents were stirred to ensure homogeneous distribution of the surfactant. The cells were incubated for 1 h at 37 °C in the presence of 5% CO2. After incubation, the medium was removed and replaced with a PBS (physiological salt) solution to permit observation of the cells and qualitative assessment of the surfactant effects. The cells were observed under an inverted Zeiss Axiovert microscope, and the morphology of the cells adhered to the bottom of the wells or another vessel used. The photos of the cells were analysed with the use of the program AxioVision Release 4.6. After the observations and recording of the cell images, the solution of physiological salt was removed and replaced by an appropriate complete medium. The cells in the complete medium were left to stand for 24 h or 48 h in the incubator. Then, the cell observation was repeated, and a qualitative assessment of the surfactant cytotoxicity was made.

### Quantitative assessment of surfactant toxicity – MTT tests

The quantitative assessment of surfactant toxicity was performed on model HeLa cells and fibroblast cells from the selected cell line GM04033. Samples containing 100 μl of freshly made DMEM or EMEM medium with the abovementioned cells were supplemented with 1 μl of a water solution of a given surfactant in a series of dilutions. After surfactant addition, the contents were gently stirred. Cells were treated when the confluence reached 70% –80%. After incubation for 1 h at 37°C in an atmosphere of 5% CO_2_, the medium was replaced by complete medium containing 5% FBS and a mixture of antibiotics and antimycotic. After subsequent3 h, an MTT test was performed (Cayman, catalogue number: M2128) to establish the number of living cells. Measurements were performed using a Tecan Infinite M200Pro microplate reader (Mannedorf, Switzerland) according to the manufacturer instructions.

### Transfection efficiency of fluorescently labeled oligodeoxynucleotide

For transfection studies, similar to the toxicity studies, the culture medium was EMEM without FBS and antibiotics. The fibroblast cells of line GM03044 were approximately 70% confluent prior to the experiment. The fluorescence labelled (FITC) ssDNA (of the sequence given in Table 1) chosen for the study was synthesized by the Future Synthesis company (Poznań, Poland). The studied surfactant/ssDNA lipoplexes were investigated 20 min after the beginning of complexation. For selected abovementioned complexes, taking into account the cytotoxicity of studied surfactants and the ability to form complexes, 1 μl of lipoplex solution was added to 1 ml of the medium with fibroblast cells. Observations of the cells after transfection were made after approximately 1 h. The effectiveness of transfection was verified on the selected complex of LA14 surfactant (40 mM) and labelled ssDNA (100 μM). Again, a portion of 1 μl of the lipoplex solution was added to 1000 μl of the medium, and then the effectiveness of transfection was monitored after 20, 30, 40, 50 and 60 min. Final concentration of LA14 and ssDNA was 40μM and 100nM, respectively. The observations were made using a Zeiss Axiovert fluorescence microscope. The culture medium was replaced by PBS to reduce fluorescence of culture medium.

## Results

### Complexation and conformational changes of surfactants with dsDNA

The conformational changes in the structure of the studied nucleic acid (ssDNA or dsDNA) induced by the increase in the surfactant concentration were revealed using the circular dichroism (CD) method. Prior to the CD measurements, every lipoplex sample was analysed using agarose gel electrophoresis to evaluate the degree of complexation of DNA with lactose-based surfactants. The CD spectra obtained for the native dsDNA (comprising 21 base pairs) and a series of studied complexes with LA12 and LA14 surfactants are shown in Fig 2.

**Fig 2.**
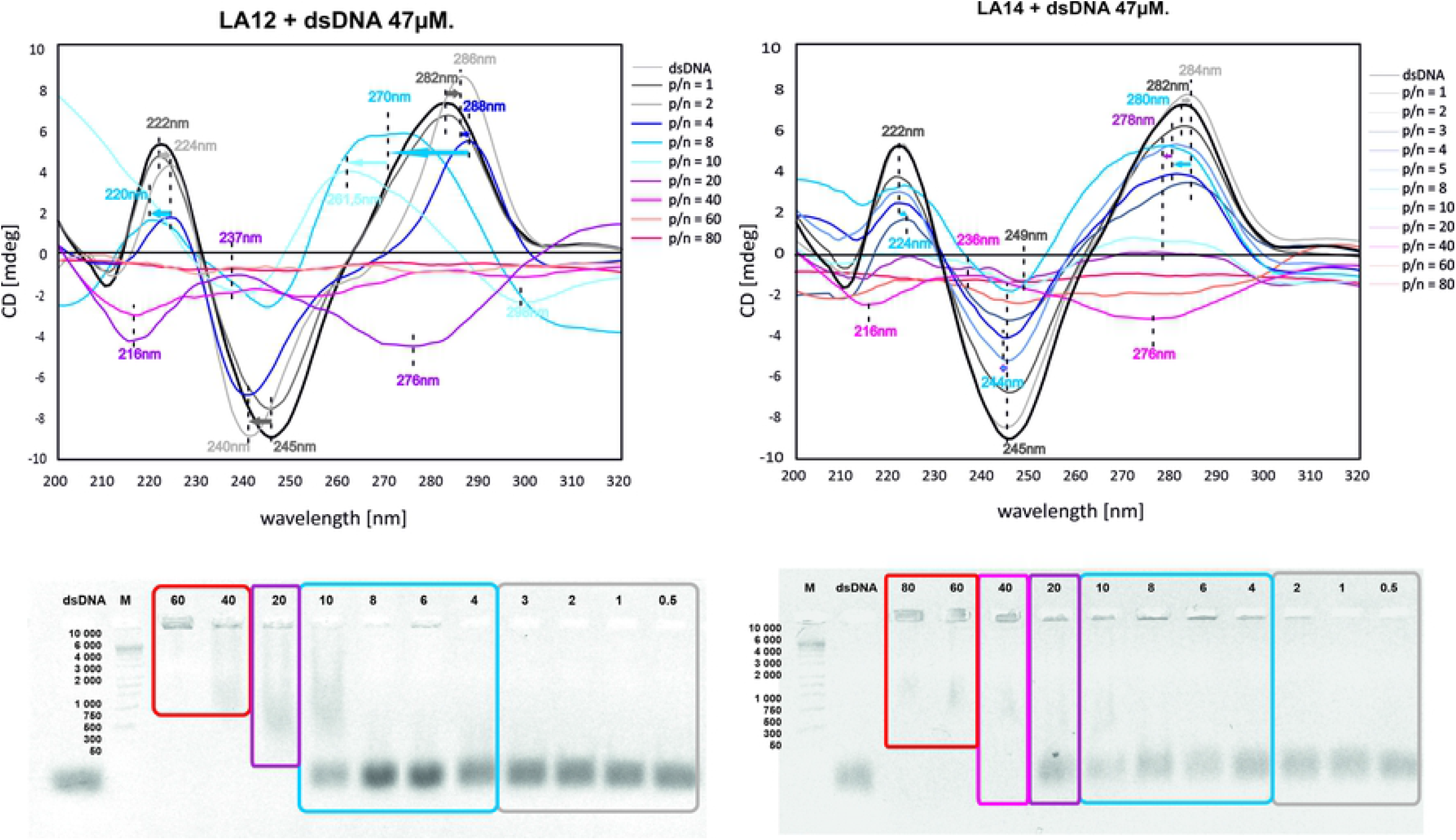
CD spectra (top) and electrophoresis results (bottom) for the dsDNA complexes with LA12 (a) and LA14 (b) surfactants. The colours of the frames correspond to those used for CD spectra, indicating the various p/n ratio ranges, whose value was determined for the individual lipoplexes over the corresponding wells. Nucleic acid (dsDNA) without surfactant was designated as control, M is the marker used.

The spectrum registered for the noncomplexed dsDNA has a positive band maximum at 282 nm and a negative band minimum at 245 nm, which are due to the positive and negative Cotton effect associated with interactions of the nearest neighbours of chromophores in the nucleic acid structure. The positions of the observed bands are characteristic of the B-DNA conformation; however, they are slightly shifted compared to those reported by Sprecher et al. [26,27]. At the lowest concentration (p/n=1, where p/n has been defined according to the equation 1), the studied surfactants do not induce significant conformational changes in double-stranded DNA. An increase in the surfactant concentration (up to p/n = 4) causes a gradual decrease in the intensity of peaks, which is accompanied by their slight shift towards longer wavelengths.

For the LA12 lipoplex (Fig 2a) at p/n = 8, the maximum (appearing at 270 nm) was significantly shifted towards shorter wavelengths, while the location of the minimum (at 244 nm) was changed only slightly compared to those observed at lower concentrations. Similar changes in the DNA spectrum have been reported earlier [28,29] and recognized as being due to the formation of Z-DNA. Further changes in the CD spectrum observed for p/n = 20,manifested by the reversed sequence of positive and negative bands, correspond to the formation of the levorotatory DNA helix (possibly of the Z-DNA form [28]); however, the CD spectra extremes are slightly shifted towards shorter wavelengths. The gel electrophoresis pattern confirms a partial complexation for this LA12 lipoplex (Fig 2a). A further increase in the surfactant concentration in the complex leads to gradual disappearance of the positive and negative bands in the CD spectrum, up to their total flattening at p/n = 50. The disappearance of the bands is accompanied by an increase in the opacity of the sample. The gel electrophoresis pattern indicates complete complexation for p/n above 40.

For the LA14/dsDNA complex at a concentration of p/n = 8, there is a visible shift of the maximum of the CD spectrum (Fig 2b), but it is not as large as for LA12. An increase in concentrations of p/n above 10 leads to a strong, progressive reduction of the intensities of the bands, which may be assigned to the hydration of the DNA oligomer helix in the vicinity of phosphate groups and sugars. A further increase in concentration leads to changes in the lipoplex CD spectrum, which at p/n = 40 is inverted, has a low intensity and is slightly shifted towards shorter wavelengths compared to that characteristic of the B-DNA form. A very similar CD spectrum, although of higher intensity, was observed for the LA12/dsDNA complex at twice lower the concentration and retarded electrophoretic mobility. With a further increase in the surfactant concentration in the complex, the positive and negative bands in the CD spectrum gradually disappear until complete flattening is observed for p/n = 80.

### Complexation and conformational changes of surfactants with ssDNA

The CD spectra obtained for native ssDNA (consisting of 23 nucleotides) and its complexes with LA12 and LA14 are shown in Fig 3.

**Fig 3.**
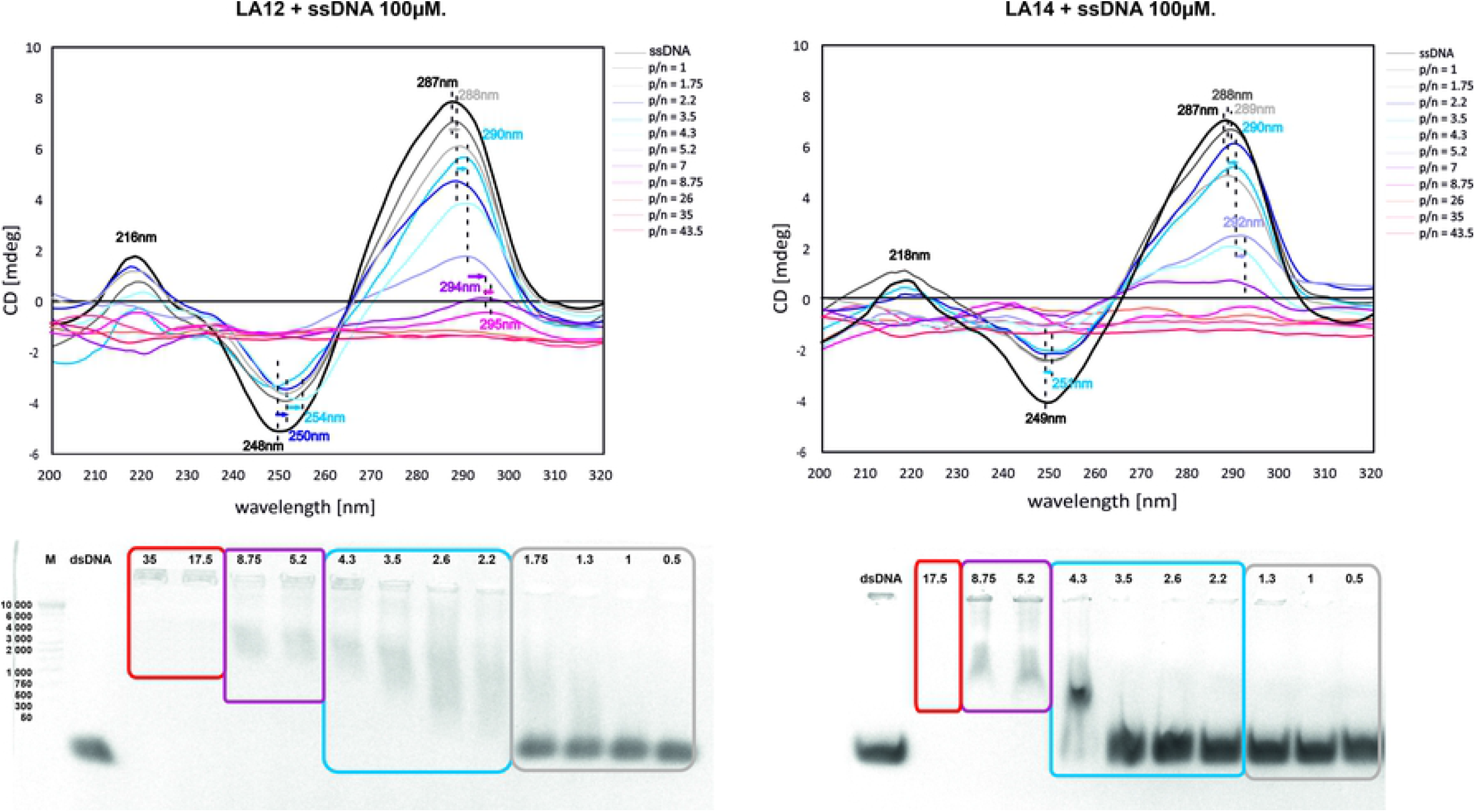
Comparison of CD spectra and electrophoresis results for ssDNA complexes with LA12 (a) and LA14 (b). The colours of the frames correspond to those used for CD spectra, indicating various p/n ratio ranges, whose value was determined for the individual lipoplexes over the corresponding wells. Nucleic acid (ssDNA) without surfactant was designated as control, M is the marker used.

The spectrum of the uncomplexed single-stranded DNA differs significantly from that observed for double-stranded DNA forms, namely, it consists of two positive peaks: an intensive peak at 216 nm and a much lower peak at 287 nm and one negative peak at 248 nm. The CD spectrum of this ssDNA structure shows a shift of the positive band towards longer wavelengths with increasing surfactant concentration. At the lowest concentrations (for p/n lower than 1.3), surfactants do not significantly affect the shape of the CD spectrum of the lipoplex; only a slight shift of the positive band towards longer wavelengths is observed. With an increase in p/n (above 1.3), both the intense positive and negative bands in the CD spectrum shift towards longer wavelengths, and their intensities decrease with increasing surfactant concentration. In contrast, a further increase in p/n (at 3.5 and 4.3) causes a rapid increase in the intensity of the bands, which is accompanied by their further shift towards longer wavelengths. At p/n = 5.2, the spectrum consists of a positive band at 290 nm and a wide negative band extending up to 200 nm. For the two subsequent surfactant concentrations (p/n = 7 and 8.75), a gradual flattening of the intensive positive band (at 290 nm) and disappearance of the weak positive band occur. These changes correspond to the increasing contributions of the chromophore interactions with the second and third neighbours. A further increase in the surfactant concentration results in loss of the CD signal, which may be due to the sample opacity.

### Structural changes in lipoplexes with DNA

Complementary to the abovementioned structural investigations were synchrotron radiation small angle X-ray scattering (SR-SAXS) studies, which revealed the structural and associative changes taking place in the solutions of the studied lipoplexes (with dsDNA or ssDNA) as a result of an increase in surfactant concentration (supplementary data: S1 Table and S2 Fig). The SR-SAXS scattering curves obtained for complexes of double-stranded DNA (dsDNA) with LA12 and LA14, as well as for pure nucleic acid solution, are shown in Fig 4.

**Fig 4.**
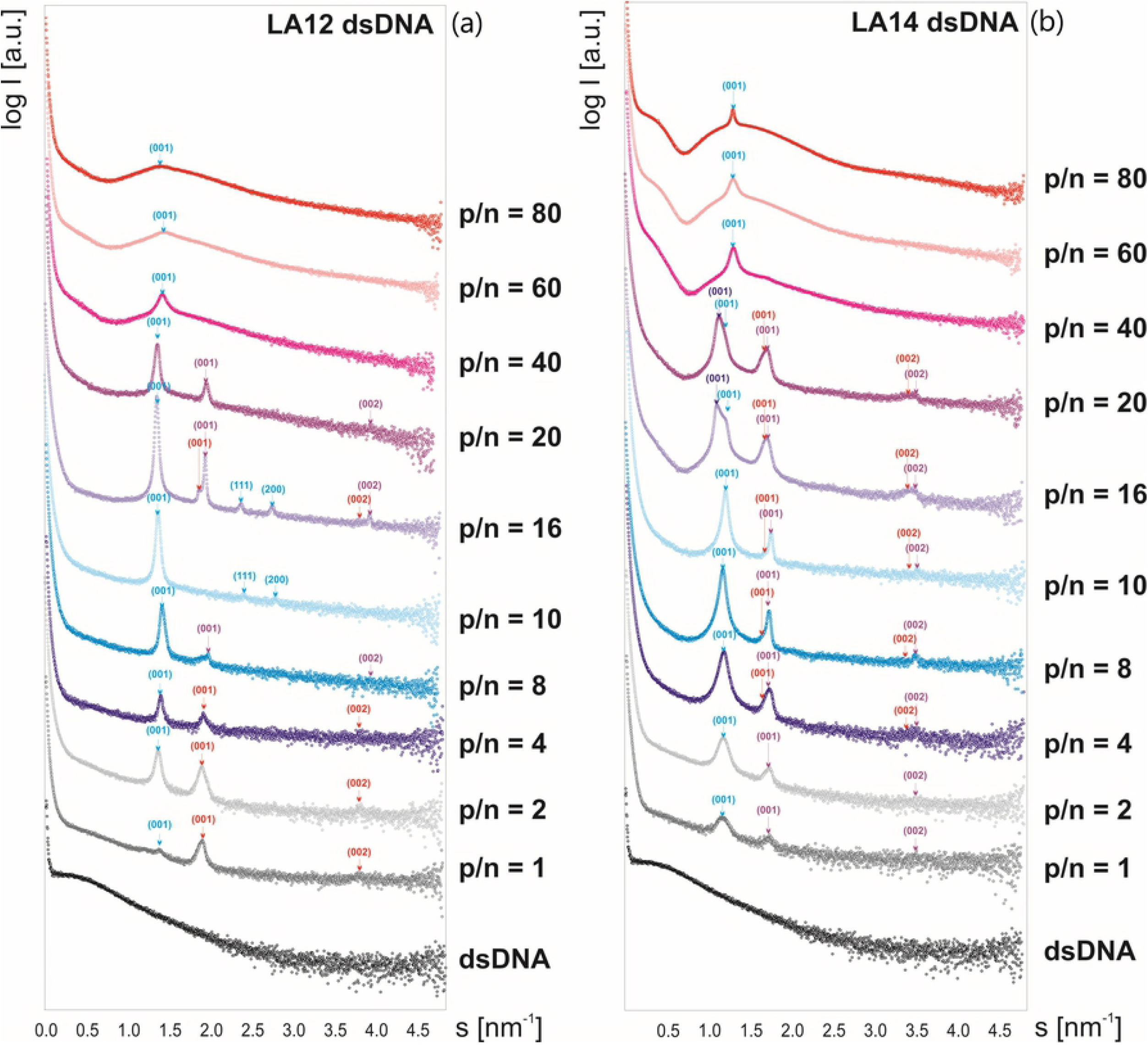
SAXS curves obtained for dsDNA complexes with (a) LA12 and (b) LA14. The hkl indices are assigned to the hexagonal structure (blue) and lamellar structures of the systems studied (red and violet).

The results of our previous SAXS studies for pure LA12 solutions [8] were taken into account in the analysis of the phase structure of the studied dsDNA lipoplexes (at various surfactant concentrations). The SR-SAXS curves obtained for the lipoplexes of the dsDNA with the LA12 surfactant (of shorter alkyl chains, Fig 4a) at the lowest concentrations (p/n below 4) show the maxima, which may be assigned to the single lamellar phase (d_001_= 3.29 nm) and to the hexagonal structure formed in the system (the dominant peak corresponding to d_100_= 4.52 nm and a_0_= 5.22 nm). Such single lamellar phase has also been revealed for the pure LA12 solution [8]. For the system characterized by p/n equal to 10 (corresponding to 20 mM LA12),the lamellar phases disappear, but the peaks characteristic of the hexagonal phase remain (d_100_= 4.42 nm, a_0_= 5.1 nm). With a further increase in the surfactant concentration (for p/n = 16 and 20), the lattice constants increase to a_0_values of 5.18 nm and 5.26 nm, respectively, which is accompanied by the reappearance of both coexisting lamellar phases (d_001_= 3.29 nm and d_001_= 3.19 nm). At higher surfactant concentrations (p/n above 40), the shape of the SAXS curves proves the dominance of liposomal structures.

At the smallest concentrations of LA14 in the system with dsDNA (Fig 4b), the SAXS curve indicates the existence of a single lamellar phase (d_001_= 3.59 nm) as well as another, which, due to the similarity of the data obtained for both surfactant systems, was assumed to be the hexagonal phase (d_100_= 5.23 nm, a_0_= 6.05 nm). Further growth in the LA14 concentration in the complex results in a change in the lattice parameter of the lamellar phases (d_100_= 3.59–3.61 nm and d_100_= 3.69–3.74 nm). Moreover, at higher concentrations of this surfactant (32 mM and 40 mM, which correspond to p/n = 16 and 20), two hexagonal phases coexisted. The lattice parameter a_H1_ for the first one is 5.8 nm, while for the second one, a_H2_ changes with increasing concentration from 6.10 nm to 6.37 nm at 32 mM and 40 mM, respectively. At the highest LA14 concentrations, the shape of the SAXS curves indicates the dominance of liposomal structures.

The SR-SAXS scattering curves (shown Fig 5) were obtained for complexes of single-stranded DNA (ssDNA) with LA12 and LA14 surfactants, as well as for the pure nucleic acid solution. The SR-SAXS scattering curves obtained for complexes of single-stranded DNA (ssDNA) with two studied surfactants show similar dependencies. Above a concentration of p/n = 8 (surfactant concentration of ca. 20 mM), lipoplexes with ssDNA appear in the aforementioned lamellar phases, characteristic of the individual surfactant solutions. For complexes with LA14, the lamellar structure is present up to p/n = 20, and the lattice constant of this phase (d_001_= 3.74 nm) is 0.54 nm larger than in the systems with LA12 (d_001_= 3.2 nm). The shape of the SAXS curves for the studied lipoplexes at the highest concentrations indicates the formation of a stable complex of ssDNA with each of these surfactants. In the systems of ssDNA complexed with the surfactants used at concentrations corresponding to p/n higher than 8.75, the appearance of a lamellar phase coexisting with the micellar phase was observed. At higher surfactant concentrations (p/n above 20), the shape of the SAXS curves suggests the dominance of liposomal structures, similar to the surfactant/dsDNA systems.

**Fig 5.**
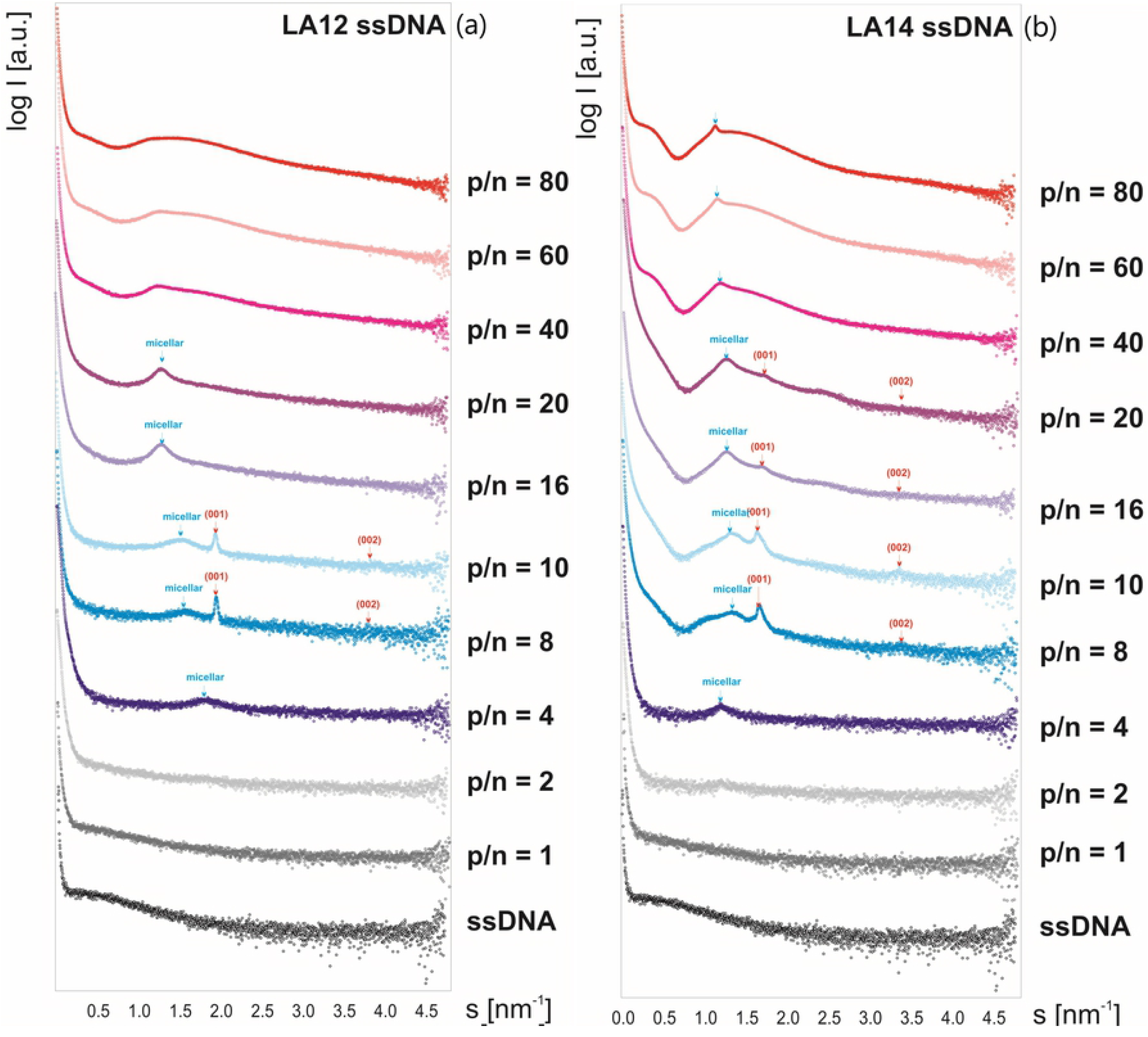
SAXS curves for ssDNA in complex with (a) LA12 or (b) LA14. Red (hkl) indices correspond to the lamellar structures of the systems.

### Complexation and conformational changes in siRNA/surfactant systems

Changes in the conformation of double-stranded RNA (siRNA) induced by the presence of the LA14 surfactant in the system were found on the basis of analysis of the CD spectra obtained for their complexes (see Fig 6). Our earlier studies on complexation in the siRNA/LA12 system [8] were taken into account in the interpretation of the obtained results.

**Fig 6.**
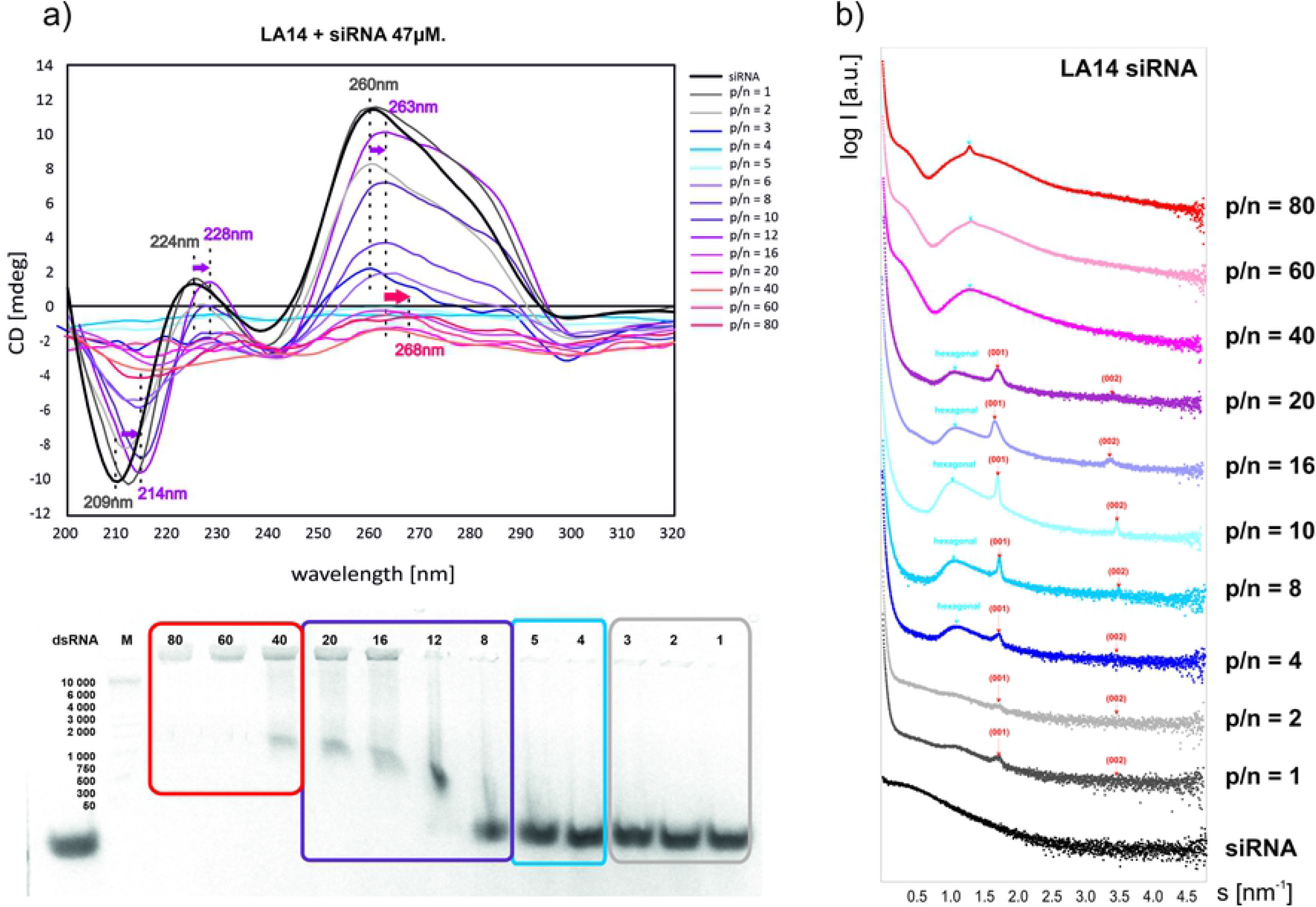
(a) CD spectrum and results of electrophoresis for the complexes of siRNA with LA14.The colours of the frames correspond to those used for CD spectra, indicating various p/n ratio ranges, whose value was determined for the individual lipoplexes over the corresponding wells. Nucleic acid (dsRNA) without surfactant was designated as control, M is the marker used. **(b) SAXS curves obtained for siRNA complexes with LA14.** The hkl indices (red) correspond to the lamellar phases formed in the systems studied.

For the complexes of siRNA with the LA14 surfactant (with a longer alkyl chain compared to LA12), the changes observed in the CD spectrum and electrophoretic pattern (Fig 6a) are similar to those observed for siRNA complexes with LA12 [8]. Taking into account the results obtained for LA12 and its complexes with siRNA, it was found that the A-RNA form of the nucleic acid is formed in the complex with LA14 at 40 mM surfactant concentration. For lower concentrations (p/n below 4), a gradual decrease in CD spectrum intensity without significant shape changes was observed. The further increase in the LA14 surfactant concentration (p/n above 4) leads to a flattening of the CD spectrum, which is a consequence of the turbidity of the solution. Interestingly, for p/n of value 6, the reconstruction of the spectrum is discerned, while the main maximum (260 nm) has been shifted towards higher wavelengths. For higher concentrations, there is a visible decrease in the intensity of the CD spectrum, which flattens above p/n equal to 20. The lack of electrophoretic mobility revealed at the highest studied surfactant concentrations indicates that siRNA is then fully complexed.

In our previous work, we showed that for the lowest three concentrations of LA12 (p/n = 1, 2 or 4) in the system with siRNA, a single lamellar phase is formed (d_001_= 3.17 nm), which is also observed for pure surfactant and for the systems of LA12 with dsDNA (Fig 4a). In the systems with higher LA12 concentrations (p/n = 8, 10 or 16), the presence of coexisting lamellar phases of d_001_=3.28 nm and d_001_=3.17 nm is observed, both for pure surfactant and its complexes with DNA at the same concentrations^8^. The shape of SAXS curves obtained for the siRNA complexes with LA14 (Fig 6b) does not correspond to that observed for LA12 systems. For these complexes, at p/n values from 4 to 20, the formation of a single lamellar phase characterized by d_001_=3.75 nm is observed (also detected in the complexes with dsDNA). Moreover, for these systems, a broad peak in the SAXS curves, probably originating in the forming hexagonal structure, reappears with increasing concentration. The SAXS curves of the systems at higher concentrations show the broad diffraction maximum corresponding to the lattice parameter of d_100_= 4.69 nm (a_0_= 5.42 nm) and thus of the values characterizing the hexagonal structures formed in the abovementioned systems with dsDNA (Fig 4b).

### Cytotoxicity of the studied surfactants

We predict, that studied surfactant may show some cell toxicity in higher concentration. Therefore, to see whether they can be used as potential tranfection reagents we first tested their effect on growing cells using microscopic and cell viability tests. the results of qualitative rough evaluation of the number and morphology of cells treated with different concentration of LA14 is shown on Fig 7. Cytotoxicity of LA12 has already been assessed by Skupin et al.[8].

**Fig 7.**
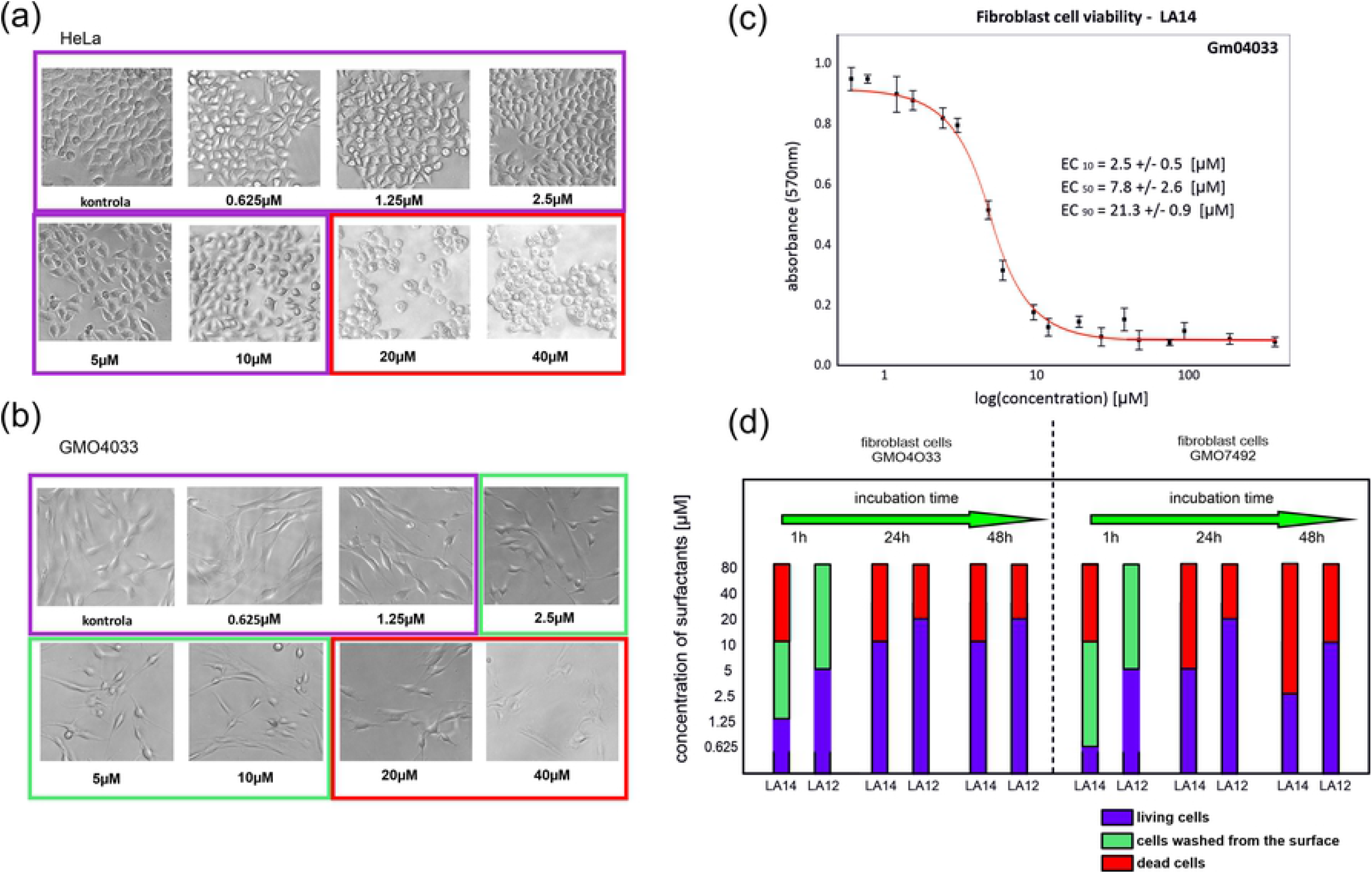
Microscopic photographs of cells surrounded by molecules of surfactant LA14 after 1 h of incubation for the three cell lines (a) HeLa, (b) GM04033, and (c) GM07492 (d) MTT test on GM04033 cell line. Purple colour is used for living cells, green colour is for cells washed from the surface, and red is for dead cells.

According to the microscopic observations performed after 1 h of incubation (Fig 7a) under specified conditions (see Materials and methods section), the addition of LA14 at concentrations from 0.625 μM to 10 μM to HeLa cells does not lead to significant changes in the morphology and number of the cells. A reduced number of cells and significant morphological changes in the cells were observed for LA12 added to HeLa cells at a concentration of 40 μM [8], while analogous changes in the systems with LA14 were noted at a concentration of 20 μM. Observations of HeLa cells in the presence of LA14 at concentrations of 20 μM and 40 μM revealed high mortality of the cells manifested as their breaking from the substrate, the appearance of grains and destruction of cell membranes (the images framed in red in Figure 7a, b and c). For the incubation time close to 24 h or 48h, no morphological changes and a high confluence of cells, comparable to that observed for the reference sample, are noted for surfactant concentrations from 0.625 μM to 20 μM.

In the tests with the GM04033 and GM07492 fibroblast lines, after 1 h of incubation, significant morphological changes were observed for certain concentrations of the surfactant. For LA14 at concentrations from 0.625 μM to 2.5μM, no morphological changes were noted. For LA14 at concentrations from 2.5 μM to 20 μM, the number of cells breaking off from the substrate increased with increasing LA14 concentration (see the images framed in green in Figures 7b and c). In the presence of LA14 at concentrations higher than 2.5 μM, the cell shape becomes more rounded, and cells break off from the substrate; however, no other significant morphological changes, such as granularity or cell membrane damage, are observed. The cells seem to be washed out of the surface. The next stage of the study was aimed at quantitatively confirming the qualitative results concerning the toxicity of the studied surfactants. The applied method was the MTT colorimetric assay, which permits quantitative determination of metabolically active cells in the tested population. On the basis of the test results, the effective surfactant concentration EC50, which corresponds to the LA14 concentration in which 50% of cells remain on culture dish compare to the control sample. was estimated. The cytotoxicity of LA14 towards HeLa cells and GM04033 fibroblasts is illustrated by the MTT absorbance curves and EC_50_ values (see S1 Fig. in supporting information and Fig 7c).

Comparison of the earlier reported cytotoxicity data for LA12 [8] with the results obtained for LA14 confirms the lower cytotoxicity of LA12 and its lower impact on HeLa cells than on fibroblasts. The effective concentration EC50 describing the impact of a surfactant solution on HeLa cells was 52.2 μM for LA12 and 23 μM for LA14. The impact of the surfactants on GM04033 fibroblasts is described by lower EC_50_ values of 14.4μM for LA12 and 7.8μM for LA14. In further transfection tests, surfactants were used at concentrations ranging from the above-determined values to take into account the different toxicities of pure surfactants and their complexes with nucleic acids.

### Transfection of fluorescently labelled ssDNA complexed with LA14

The transfection tests were performed on GM04033 cells for the LA14/ssDNA complex using ssDNA labelled at 5’-endwith a fluorescent FITC dye. The first experiment, aimed at checking the relative effectiveness of transfection, was performed for the complex in which LA14 had a concentration of 40 mM and ssDNA had a concentration of 100 μM (p/n = 17.5). On the basis of the electrophoretic mobility tests (arrest of mobility of nucleic acid) and the observations of flattening of the CD spectrum of the system surfactant and DNA (corresponding to appearance of turbidity of the sample and formation of a stable complex), as it was found, LA14 at this concentration is most effective in complex formation with DNA. The effectiveness of transfection was established on the basis of monitoring the sample studied at 10 min intervals starting from addition of the complex to the culture dish with the cells. Fifty min after the addition of the complex, the transfection was clearly visible (Fig 8a).

**Fig 8.**
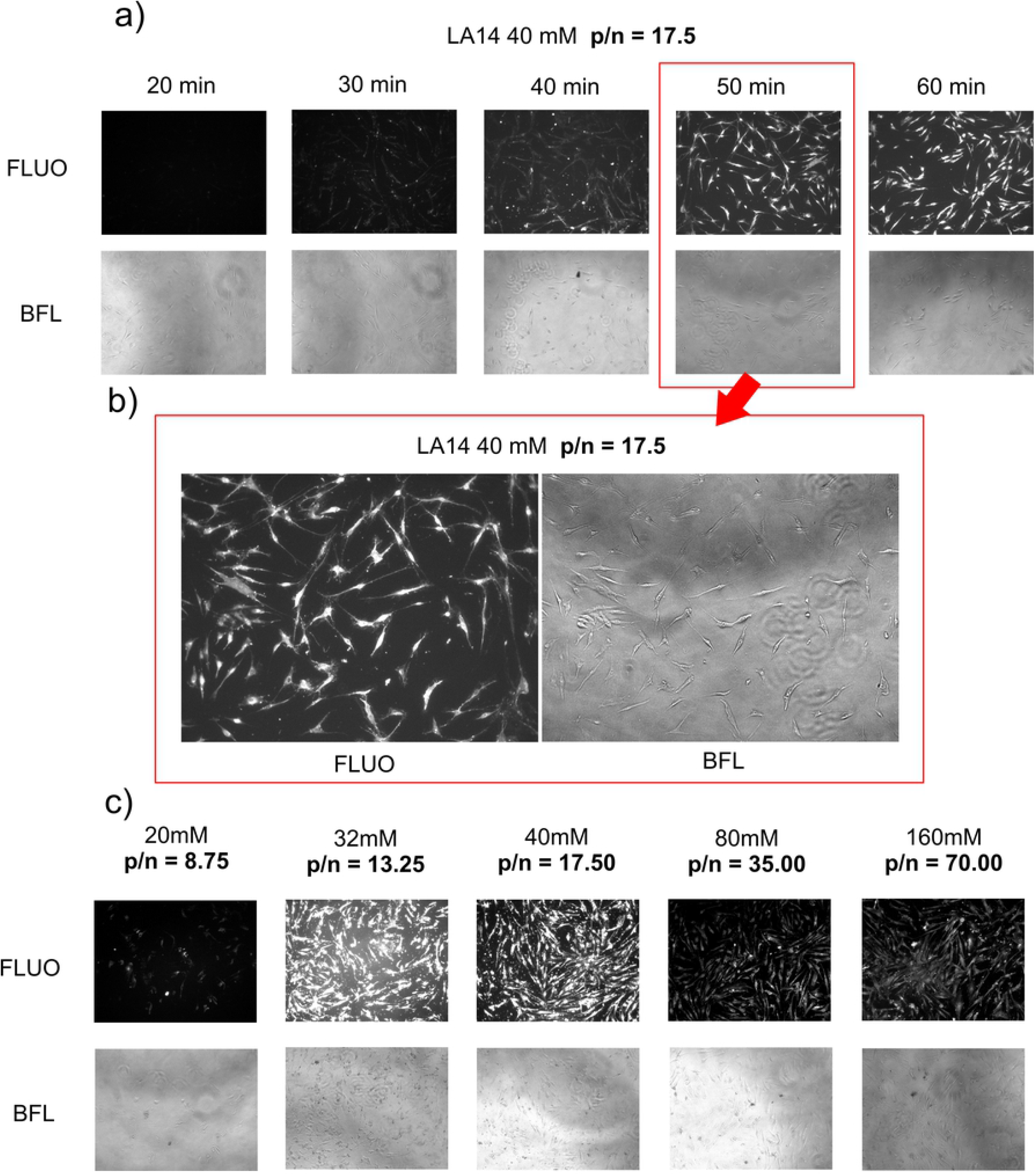
(a) Visualization of the transfection process of the LA14/ssDNA complex with LA14 at a concentration of 40 mM and ssDNA at 100 μM (p/n=17.5) for the indicated transfection times. (b) Enlarged images obtained after 50 min of transfection.(c) Visualization of the transfection for LA14/ssDNA complexes at various concentrations of the LA14 surfactant. Fluorescence microscopy images show FITCssDNA (FITC – fluorescein isothiocyanate) penetration into cells labelled FLUO, while bright-field light images show BFL.

Thus, in further studies, the effectiveness of transfection was checked 60 min after the addition of the complex, because this time was optimal for realization of transfection. In the next step, to identify the most effective complex of ssDNA with LA14, transfection was performed using a transfection solution obtained by mixing different concentrations of surfactant from 20mM to 160mM and the same 100μM concentration of ssDNA (Fig 8b). The best conditions of transfection were established on the basis of the fluorescence from the ssDNA transfected to the cells and the cell viability. The highest fluorescence intensity and the greatest cell viability were noted for ssDNA/LA14 complexes at two concentrations, i.e., 32 mM and 40 mM (corresponding to the ratios p/n = 13.25 and 17.5, respectively).

## Discussion

Based on the analysis of the CD spectra for the surfactant/dsDNA complexes in the low concentration surfactant range (below p/n = 4), the B-DNA conformation was found, and only its slight changes with the surfactant concentration increase were detected for both LA12 and LA14. These changes may be explained as resulting from the interactions between the positively charged surfactant groups and the polyanionic DNA molecule^30^. Presumably, due to the interaction between dsDNA and surfactant molecules, the hydrophobic part of the surfactant becomes exposed to the solution, which leads to an increase in the local environment polarity of the chromophores, causing local disturbances in the geometry of bases in the DNA helix. A further increase in surfactant concentration leads to a change from a right-handed helix conformation (B-DNA) to a left-handed conformation, the most similar to the Z-DNA form. However, the formation of the Z-DNA structure of the duplex cannot be unambiguously confirmed because of the low intensity of the spectrum; however, based on the observations of changes in the CD spectrum (Fig 2a), it can be concluded that the geometry of nucleic bases is similar to that of the Z-DNA form, revealing a shift in the CD spectra extremes towards shorter wavelengths as a consequence of the changes in polarity of the chromophore environment and exposition of the hydrophilic components of the surfactants towards the solution. This effect is accompanied by partial complexation of the lipoplexes, which was revealed using electrophoretic tests. For the highest concentrations used, the lack of electrophoretic mobility of DNA in the agarose gel was observed, indicating that the negative charge of DNA is completely neutralized by interacting surfactant molecules. Moreover, it was found that both conformational changes and complexation processes occur at lower concentrations for surfactant LA12 (with shorter chains) in comparison with LA14.

The small-angle X-ray scattering method was used to correlate the conformational changes of dsDNA within lipoplexes and the spatial arrangement of surfactants in aqueous solution. The lamellar phase was revealed from SAXS data of aqueous solutions of both LA12 and LA14 surfactants. The changes in SR-SAXS curves for LA12/dsDNA lipoplexes are similar to those discerned for surfactants with longer alkyl chains (LA14). The determined difference between the d-spacings for these structures corresponds to the difference between the alkyl chain lengths of these two surfactants (the length of the CH_2_-CH_2_ unit). The presence of dsDNA in the system causes the appearance of a hexagonal phase in addition to the abovementioned lamellar phase. The surfactant concentrations at which the hexagonal structure becomes dominant correlate well with the observations of the initiation of conformational changes into the Z-DNA form disclosed in CD spectra as well as the loss of electrophoretic mobility of dsDNA complexed with the surfactant. At the highest surfactant concentrations, the shape of the SAXS curves indicates the dominance of liposomal structures.

The ssDNA has a helical structure, similar to dsDNA, which implies a regular distribution of chromophores and a significant impact of interactions between the nearest neighbours on the shape of the CD spectrum. The spectrum of the uncomplexed singlestranded DNA differs significantly from that observed for double-stranded DNA forms. For low surfactant concentrations, the shape of the curves remains unchanged, but a slight shift of the positive band of ssDNA towards longer wavelengths (observed at p/n higher than 3.5) results from interactions with surfactants, similar to what was observed for dsDNA. The disappearance of the main positive band observed for p/n higher than 8.75 (surfactant concentration above 20 mM) suggests compensation of the interactions between the nearest neighbours. The results of electrophoretic tests indicate the formation of stable complexes of surfactants with the nucleic acid.

The small-angle X-ray scattering method was used to correlate the conformational changes of ssDNA within lipoplexes and the spatial arrangement of surfactants in aqueous solution. A difference of 0.54 nm between the values of these constants in the LA12/ssDNA and LA14/ssDNA systems corresponds to the doubled difference in length of chains of these surfactants (four CH_2_ segments), suggesting the formation of double lamellar structures in the complex with the nucleic acid. The formation of a stable complex of ssDNA with each of these surfactants correlates well withthe observations of the CD spectra of the complexes, which indicate a significant conformational change of ssDNA (flattening of the CD signal) at p/n above 8.75 as well as electrophoretic mobility tests evidencing the lack of mobility of such complexes.

It was found that the LA14 surfactant induces the same conformational changes in siRNA as LA12, although at much lower concentrations.

The CD spectra obtained for LA14/siRNA systems indicate the presence of the A-RNA structure in these systems. The increase in the surfactant concentration results in flattening of the signal, most likely due to the interaction of the surfactant with individual bases of the nucleic acid. The electrophoretic mobility suggests the lack of stable complex formation at these concentrations. Above p/n = 6, we observe are storation of the CD spectrum for siRNA, which is associated with a change in the turbidity of the sample, most likely resulting from a different spatial arrangement of the surfactant, which affects the turbidity of the solution. At p/n=20 (40 mM surfactant concentration), the CD signal of RNA is flattened, and electrophoretic tests prove the loss of mobility of the lipoplexes, which confirms the formation of stable complexes in the system.

Structural analysis of siRNA/LA12 lipoplexes and the pure surfactant solutions at various concentrations, based on SAXS data, have been reported in our previous work^8^. However, taking into account the results of the abovementioned SAXS studies of systems with various nucleic acids and the tendency of the tested systems to adopt a certain packing, it is possible to propose an alternative interpretation of results obtained for the highest LA12 concentrations in the systems with siRNA.

In the systems with siRNA, the formation of lamellar structures was also observed; for LA12, two coexisting phases were characterized by d-spacing values of d=3.28 nm and d=3.17 nm, while for LA14, only one phase was characterized by d=3.57 nm. Similar to the systems with DNA, these d-spacing values probably correspond to the structures formed by surfactants complexed with siRNA. The scattering curves of the siRNA/LA12 systems show strong peaks corresponding to the hexagonal structure (d=4.52 nm; a_0_=5.22 nm).The difference in the structural parameters observed for LA12 complexes with RNA and DNA is a consequence of a small difference in the diameter of these two nucleic acid molecules. Moreover, for these systems (with the same p/n ratios), a characteristic broad peak appears, which has been interpreted in our previous work as corresponding to the micellar phase of the surfactant. However, taking into account the formation of hexagonal phases discussed in this paper systems with dsDNA, it is reasonable to assume that this additional broad peak could be ascribed to the hexagonal phase. Additionally, with the appearance of the peak from the hexagonal phase, we observe the reconstruction of the A-RNA spectrum and the lack of turbidity of the solution, which confirms the structural change taking place for the complex.Our previous interpretation [8] assumed that the complexes with LA12 at the highest concentrations (p/n = 20, 40 or 80) were from the regular structure Im3m and the coexisting lamellar structure. In view of the results obtained within this work, including SAXS results and spectroscopic data obtained for different systems based on LA12 and LA14 surfactants, it can now reasonable be assumed that instead the cubic phase, identified on the basis of a few diffraction maxima, could also be a combination of the coexisting lamellar phases and a hexagonal phase of the size that is close to that obtained for other systems with the same surfactant. In the siRNA/LA12 systems, the lamellar phases are characterized by d_001_= 3.28 nm and d_001_=3.17 nm, while the hexagonal phase is characterized by d_100_= 4.52 nm and a = 5.22 nm, which are the same as the parameters of the structures observed for the LA12 complexes with dsDNA (see Fig 4). For the systems with the LA14 surfactant at higher concentrations, the hexagonal phase is not directly revealed, as was found for the systems with LA12 [8], but the SAXS data indicate the formation of large liposomal forms, the presence of which could have obscured or dominated the hexagonal form.

According to the analysis of the results concerning the toxicity of the studied surfactants, all cell lines used in the experiment show higher tolerance to the surfactant with the shorter alkyl chain (LA12). Both surfactants were also noted to wash out the cells from the surface of culture dish, but without changes in their morphology. This observation indicates a very strong reduction in the surface tension and possible interactions of surfactant molecules with the compounds on the cell membranes. For fibroblasts, i.e., cells with less tolerance to external factors, as well as HeLa cells, morphological changes of cells induced by the presence of surfactants were observed at different surfactant concentrations and after different incubation times. For the surfactant with the shorter alkyl chain (LA12) [8], the mean surfactant concentrations for cell survival ranged from 10 μM to 20 μM, while for surfactant with a longer side chain (LA14), the mean surfactant concentrations ranged from 2.5 μM to 10 μM. Above these concentrations of surfactants, the cells are observed to be washed out from the surface, which, however, is not tantamount to their death. The results of the transfection experiment allowed to determine the effectiveness of this process, taking place in approximately 50 min from adding complex mixture, and select the most effective transfection system for the ssDNA-FITC/LA14 complex, which corresponded to a p/n ratio higher than 8.75. The new surfactants based on lactose derivatives of betaines showing zwitterionic character were found to be very effective carriers of short nucleic acids and show potential for application in more advanced an complex studies in the future.

## Conclusion

The results of the research based on the series of experiments performed for the two studied surfactants LA12 and LA14, differing in the length of the alkyl chain, imply a much better ability of LA12 to form complexes with nucleic acids when the former is used at lower concentrations. In the systems of nucleic acids with either of the two surfactants, the dominant lamellar and hexagonal structures were formed. Moreover, it was found that the increase in the surfactant concentration induces conformational changes in the nucleic acids. It was proven that the LA12 surfactant (with a shorter alkyl chain) showed the best complexing ability with each of the three studied nucleic acids (siRNA, dsDNA and ssDNA) when used during complexation at a concentration of 40 mM, while for LA14, the optimal concentration was twice as high (i.e., 80 mM), except for complexes with ssDNA when LA14was most effective at 40 mM.

## Acknowledgments

This research has been founded by the Ministry of Science and Higher Education (Poland) Diamond Grant program No. 0011/DIA/2015/44 (to MW) and by National Science Centre (Poland)ETIUDA 8 DEC-2020/36/T/ST4/00099 (to MW) grant and 2020/38/A/NZ3/00498 (to KS). Michalina Wilkowska got a scholarship from the Adam Mickiewicz University Foundation for the 2019/2020 academic year.

## Disclosure

The author reports no conflicts of interest in this work.

## Supporting information

**S1 Fig. The MTT absorbance curves and EC_50_ values of the cytotoxicity of LA14 towards HeLa cells**

**S2 Fig. SAXS curves obtained for LA14 surfactant in different concentrations. S1 Table. Table summarises SAXS studies results for LA12 and LA14.**

## Notes

### Competing Interest Statement

The authors have declared no competing interest.

## References

1. Shichiri M, Tanaka A, Hirata Y. Intravenous gene therapy for familial hypercholesterolemia using ligand-facilitated transfer of a liposome:LDL receptor gene complex. Gene Ther. 2003 May;10(9):827–31. doi: 10.1038/sj.gt.3301953.

2. Cerchia L, Hamm J, Libri D, Tavitian B, de Franciscis V. Nucleic acid aptamers in cancer medicine. FEBS Lett. 2002 Sep 25;528(1-3):12–6. doi: 10.1016/s0014-5793(02)03275-1.

3. Demeunynck, M., Bailly, C. & Wilson, W.D. Small Molecule DNA and RNA Binders: From Synthesis to Nucleic Acid Complexes. John Wiley & Sons, 2006.

4. Gerson, S.L. & Lattime, E.C. Gene Therapy of Cancer: Translational Approaches from Preclinical Studies to Clinical Implementation. Academic Press, 2002.

5. Sharma VD, Lees J, Hoffman NE, Brailoiu E, Madesh M, Wunder SL, Ilies MA. Modulation of pyridinium cationic lipid-DNA complex properties by pyridinium gemini surfactants and its impact on lipoplex transfection properties. Mol Pharm. 2014 Feb 3;11(2):545–59. doi: 10.1021/mp4005035.

6. Ogris M, Walker G, Blessing T, Kircheis R, Wolschek M, Wagner E. Tumor-targeted gene therapy: strategies for the preparation of ligand-polyethylene glycol-polyethylenimine/DNA complexes. J Control Release. 2003 Aug 28;91(1-2):173–81. doi: 10.1016/s0168-3659(03)00230-x.

7. Bombelli C, Faggioli F, Luciani P, Mancini G, Sacco MG. Efficient transfection of DNA by liposomes formulated with cationic gemini amphiphiles. J Med Chem. 2005 Aug 11;48(16):5378–82. doi: 10.1021/jm050477r.

8. Skupin M., Sobczak K., Zieliński R., Kozak, M. The System with Zwitterionic Lactose-Based Surfactant for Complexation and Delivery of Small Interfering Ribonucleic acid—A Structural and Spectroscopic Study. Appl Phys Lett. 2016 108(21): 213701-1-5. doi: 10.1063/1.4952589.

9. Wojtkowiak-Szlachcic A, Taylor K, Stepniak-Konieczna E, Sznajder LJ, Mykowska A, Sroka J, Thornton CA, Sobczak K. Short antisense-locked nucleic acids (all-LNAs) correct alternative splicing abnormalities in myotonic dystrophy. Nucleic Acids Res. 2015 Mar 31;43(6):3318–31. doi: 10.1093/nar/gkv163.

10. Kotula JW, Pratico ED, Ming X, Nakagawa O, Juliano RL, Sullenger BA. Aptamer-mediated delivery of splice-switching oligonucleotides to the nuclei of cancer cells. Nucleic Acid Ther. 2012 Jun;22(3):187–95. doi: 10.1089/nat.2012.0347.

11. Catuogno S, Esposito CL. Aptamer Cell-Based Selection: Overview and Advances. Biomedicines. 2017 Aug 14;5(3):49. doi: 10.3390/biomedicines5030049.

12. Watts JK, Corey DR. Silencing disease genes in the laboratory and the clinic. J Pathol. 2012 Jan;226(2):365–79. doi: 10.1002/path.2993.

13. Dujardin G, Buratti E, Charlet-Berguerand N, Martins de Araujo M, Mbopda A, Le Jossic-Corcos C, Pagani F, Ferec C, Corcos L. CELF proteins regulate CFTR pre-mRNA splicing: essential role of the divergent domain of ETR-3. Nucleic Acids Res. 2010 Nov;38(20):7273–85. doi: 10.1093/nar/gkq573.

14. Timchenko NA, Cai ZJ, Welm AL, Reddy S, Ashizawa T, Timchenko LT. RNA CUG repeats sequester CUGBP1 and alter protein levels and activity of CUGBP1. J Biol Chem. 2001 Mar 16;276(11):7820–6. doi: 10.1074/jbc.M005960200.

15. Vlasova IA, Tahoe NM, Fan D, Larsson O, Rattenbacher B, Sternjohn JR, Vasdewani J, Karypis G, Reilly CS, Bitterman PB, Bohjanen PR. Conserved GU-rich elements mediate mRNA decay by binding to CUG-binding protein 1. Mol Cell. 2008 Feb 1;29(2):263–70. doi: 10.1016/j.molcel.2007.11.024.

16. Pietralik, Z., Taube, M., Skrzypczak, A. & Kozak, M. SAXS Study of Influence of Gemini Surfactant, 1,1’-(1,4-Butanediyl)bis 3-Cyclododecyloxymethylimidazolium Di-Chloride, on the Fully Hydrated DMPC. Acta Phys. Pol. A. 2010; 117:311–314. doi: 10.12693/APhysPolA.117.311.

17. Andrzejewska W, Wilkowska M, Peplińska B, Skrzypczak A, Kozak M. Structural characterization of transfection nanosystems based on tricationic surfactants and short double stranded oligonucleotides. Biochem Biophys Res Commun. 2019 Oct 22;518(4):706–711. doi: 10.1016/j.bbrc.2019.08.114.

18. Pietralik, Z., Krzysztoń, R., Kida, W., Andrzejewska, W. & Kozak, M. Structure and Conformational Dynamics of DMPC/Dicationic Surfactant and DMPC/Dicationic Surfactant/DNA Systems. Int. J. Mol. Sci. 14, 7642–7659 (2013).

19. Andrzejewska W, Wilkowska M, Skrzypczak A, Kozak M. Ammonium Gemini Surfactants Form Complexes with Model Oligomers of siRNA and dsDNA. Int J Mol Sci. 2019 Nov 7;20(22):5546. doi: 10.3390/ijms20225546.

20. Wojciechowska M, Taylor K, Sobczak K, Napierala M, Krzyzosiak WJ. Small molecule kinase inhibitors alleviate different molecular features of myotonic dystrophy type 1. RNA Biol. 2014;11(6):742–54. doi: 10.4161/rna.28799.

21. Kajdasz A, Niewiadomska D, Sekrecki M, Sobczak K. Distribution of alternative untranslated regions within the mRNA of the CELF1 splicing factor affects its expression. Sci Rep. 2022 Jan 7;12(1):190. doi: 10.1038/s41598-021-03901-9.

22. Blanchet CE, Spilotros A, Schwemmer F, Graewert MA, Kikhney A, Jeffries CM, Franke D, Mark D, Zengerle R, Cipriani F, Fiedler S, Roessle M, Svergun DI. Versatile sample environments and automation for biological solution X-ray scattering experiments at the P12 beamline (PETRA III, DESY). J Appl Crystallogr. 2015 Mar 12;48(Pt 2):431–443. doi: 10.1107/S160057671500254X.

23. Round A, Felisaz F, Fodinger L, Gobbo A, Huet J, Villard C, Blanchet CE, Pernot P, McSweeney S, Roessle M, Svergun DI, Cipriani F. BioSAXS Sample Changer: a robotic sample changer for rapid and reliable high-throughput X-ray solution scattering experiments. Acta Crystallogr D Biol Crystallogr. 2015 Jan 1;71(Pt 1):67–75. doi: 10.1107/S1399004714026959.

24. Huang, T.C., Toraya, H., Blanton, T.N. & Wu, Y. X-Ray Powder Diffraction Analysis of Silver Behenate, a Possible Low-Angle Diffraction Standard. J. Appl. Crystallogr. 1993; 26:180–184. doi: 10.1107/S0021889892009762.

25. Petoukhov MV, Franke D, Shkumatov AV, Tria G, Kikhney AG, Gajda M, Gorba C, Mertens HD, Konarev PV, Svergun DI. New developments in the *ATSAS* program package for small-angle scattering data analysis. J Appl Crystallogr. 2012 Mar 15;45(Pt 2):342–350. doi: 10.1107/S0021889812007662.

26. Sprecher CA, Baase WA, Johnson WC Jr. Conformation and circular dichroism of DNA. Biopolymers. 1979 Apr;18(4):1009–19. doi: 10.1002/bip.1979.360180418.

27. Zhou, T., Xu, G., Ao, M., Yang, Y. & Wang, C. DNA Compaction to Multi-Molecular DNA Condensation Induced by Cationic Imidazolium Gemini Surfactants. Colloids Surf. Physicochem. Eng. Asp. 414, 33–40 (2012).

28. Park S, Otomo H, Zheng L, Sugiyama H. Highly emissive deoxyguanosine analogue capable of direct visualization of B-Z transition. Chem Commun (Camb). 2014 Feb 14;50(13):1573–5. doi: 10.1039/c3cc48297a.

29. Kypr J, Kejnovská I, Renciuk D, Vorlícková M. Circular dichroism and conformational polymorphism of DNA. Nucleic Acids Res. 2009 Apr;37(6):1713–25. doi: 10.1093/nar/gkp026.

